# Cell state transitions drive the evolution of disease progression in B-lymphoblastic leukemia

**DOI:** 10.1101/2025.04.21.649796

**Authors:** Curtis Gravenmier, Sadegh Marzban, Yi-Han Tang, Nancy Gillis, Bijal D. Shah, Lynn Moscinski, Ling Zhang, Jeffrey West

## Abstract

Cancer stem cells (CSCs) are hypothesized to promote tumor progression through innate chemoresistance and self-renewal. CSCs reside in the CD34^+^/CD38^-^ immunophenotypic subpopulation of acute myeloid leukemia (AML); correspondingly, the size of this subpopulation has a strong negative impact on overall survival. Isolation of CSCs from B-lymphoblastic leukemia (B-ALL) has proven difficult, and the cells of interest apparently are not isolated to the CD34^+^/CD38^-^ compartment. This may be explained, in part, by temporal variations of CD34 and CD38 expression which result in stochastic cell state transitions (e.g., from CD34^+^/CD38^+^ to CD34^+^/CD38^-^). We present a mathematical model of these transitions and correlate salient findings with *BCR::ABL1* status, minimal residual disease (MRD), and relapse in adult B-ALL. As the CSC hypothesis is well supported in AML, we focus on transitions to and from the hematopoietic stem cell compartment (CD34^+^/CD38^-^), which our findings suggest can be estimated from peripheral blood or bone marrow samples. Additionally, we find that *BCR::ABL1* positive patient samples are associated with high transition rates into the CD34^+^/CD38^-^ compartment, including self-renewal. In contrast, *BCR::ABL1* negative patient samples have low CD34^+^/CD38^-^ self-renewal rates and either high CD34^+^/CD38^+^ or CD34^-^/CD38^+^ incoming rates. High CD34^+^/CD38^-^ self renewal is also associated with MRD post-induction chemotherapy. We find a lack of observable changes in cell state transitions between diagnosis and relapse specimens. Our analysis provide evidence of de-differentiation transitions to a CD34+/CD38-stem cell-like immunophenotype in leukemia samples collected from B-ALL patients, and the tendency for these transitions is especially strong for B-ALL with BCR::ABL1. The modeling framework used here is a novel, useful tool to infer both prognosis and genotype from routine flow cytometry data.

**Significance Statement:** Flow cytometry characterization of B-lymphoblastic leukemia samples (diagnosis, remission, relapse) is used to parameterize a mathematical model of cell state transition rates and stratify patients at risk for post-induction chemotherapy minimal residual disease.

## Introduction

Cancer stem cells (CSCs) are hypothesized to drive tumor progression by serving as a drug-resistant reservoir with the capacity for self-renewal. The first empirical evidence to support CSCs was obtained from human acute myeloid leukemia (AML) when it was discovered that the ability to transplant AML to severe combined immune-deficient mice is restricted to leukemia cells with a CD34^+^CD38^-^ immunophenotype similar to normal hematopoietic stem cells^1^. The progeny of transplanted CD34^+^/CD38^-^ leukemia cells acquired lineage-specific markers and reproduced the French-American-British (FAB) classification of the original human AML specimen, consistent with *in vivo* differentiation. Moreover, serial transplantation of CD34^+^CD38^-^ leukemia cells demonstrated a capacity for self-renewal, offering strong support that the CD34^+^/CD38^-^ subpopulation encompasses leukemia cells with stem cell-like properties^2^. Putative AML CSCs also possess characteristics associated with multidrug resistance. For instance, there is higher ATP-binding cassette transporter expression in CD34^+^/CD38^-^ AML cells compared to CD34^+^/CD38^+^ AML cells^3^, and a subset of CD34^+^/CD38^-^ cells is able to resist venetoclax by altering what substrates are used to fuel the tricarboxylic acid cycle and oxidative phosphorylation^4^. Accordingly, a high frequency of CD34^+^/CD38^-^ cells at AML diagnosis has a strong negative impact on survival^5–7^ and the proportion of AML cells expressing stem cell markers increases between diagnosis and relapse^8^.

Isolation of CSCs from B-lymphoblastic leukemia (B-ALL) has proven difficult, perhaps because leukemia-initiating cells are not isolated to the CD34^+^/CD38^-^ compartment^9–12^. This may be due to temporal variation of CD34 and CD38 expression which results in stochastic cell state transitions (e.g., from CD34^+^/CD38^+^ to CD34^+^/CD38^-^). Indeed, clonal subcultures derived from single B-ALL cells give rise to subpopulations with disparate CD34 and CD38 expression within hours, and ultimately reproduce a heterogenous leukemia regardless of the initial CD34 and CD38 immunophenotype^13^. Monitoring the clonal subcultures using time-lapse epifluorescence microscopy reveals temporal variation of CD34 and CD38 expression by single cells prior to any cell division, confirming that temporal variation results from transient marker expression rather than generation of differentiated progeny^13^. These observations conflict with the hierarchical nature of the CSC hypothesis and support B-ALL subpopulation dynamics are more accurately described by allowing for spontaneous, reversible cell state transitions. Still, a high frequency of CD34^+^/CD38^-^ cells at diagnosis is associated with minimal residual disease (MRD) and poor prognosis in childhood B-ALL^14–16^. In addition, a CD34^+^/CD38^-^ immunophenotype is typically observed in B-ALL with *BCR::ABL1*, which constitutes a high risk subgroup of B-ALL^17^.

### Cell state transitions drive the evolution of disease progression

Our study aims to investigate the hypothesis that B-ALL cell state transitions involving CD34 and CD38 carry prognostic significance because these transitions impact the prevalence and stability of the CD34^+^/CD38^-^ cell state. Tools based on single cell transcriptomics^18,19^ and fluorescence-activated cell sorting with flow cytometry^20,21^ have begun to unravel cancer cell state dynamics in experimental systems, but these sophisticated techniques have not yet entered the clinical laboratory. We, therefore, devised an innovative method to infer cell state transition dynamics from clinical flow cytometry data. Transitions between CD34^+^/CD38^-^, CD34^+^/CD38^+^, CD34^-^ /CD38^+^, and CD34^-^/CD38^-^ B-ALL cell states were modeled as an irreducible Markov chain. The resulting transition matrix values are useful for predicting important features of B-ALL including *BCR::ABL1* status and response to induction chemotherapy.

### Mathematical models of cell plasticity

Cancer is a complex, evolutionary disease where mathematical and computational models are highly appropriate to study and predict disease dynamics^22–24^. Previous mathematical models have considered the role of plasticity in treatment-induced stemness^25^ or other non-genetic mechanisms of drug resistance^26,27^. Other approaches consider the interplay between Lamarckian, plastic induction of resistance and Darwinian, genetic selection for resistance^28,29^. Mathematical modeling has shown promise in describing the rates of clonal expansion of hematopoietic stem cell variants^30–32^ and the dependence on microenvironmental context^33^, cytokine signaling^34^, aging^35^, or hematopoietic cell transplantation^36^. Markov chain models are a common mathematical framework to simulate time dynamics of systems with a finite number of states (e.g. cell types) where spontaneous switching between states occurs at a predictable rate or likelihood (e.g. cell state plasticity)^37–40^. For example, cell states may be inferred by lineage information and rates of cell state switching estimated by fitting Markovian models to temporal measurements^41^. Similar models extend finite states to allow for a continuous range of possible plastic phenotypes^42,43^ or continuous time^44^. The temporal kinetics of AML disease dynamics have been modelled using time-sequential bulk RNA-seq data to parameterize a state-transition model^45^. A similar model uses transcriptomics data to parameterize a state-transition model that predicts response to tyrosine kinase inhibitors in chronic myeloid leukemia^46^. Similarly, here we attempt to estimate cell state transitions involving CD34 and CD38 expression using a novel dataset of B-ALL patients, along with patients in molecular remission as a control group.

## Results

A general outline of the clinical and laboratory management of adult B-ALL is provided in **Figure 1A**. Treatment begins with induction chemotherapy, after which the patient is assessed for MRD using sensitive assays including specialized flow cytometry and clonal immunoglobulin gene sequences (ClonoSEQ). Peripheral blood and bone marrow specimens can be assessed for flow cytometric and molecular genetic evidence of residual/recurrent B-ALL throughout a patient’s treatment course in order to guide therapy decisions.

**Figure 1:**
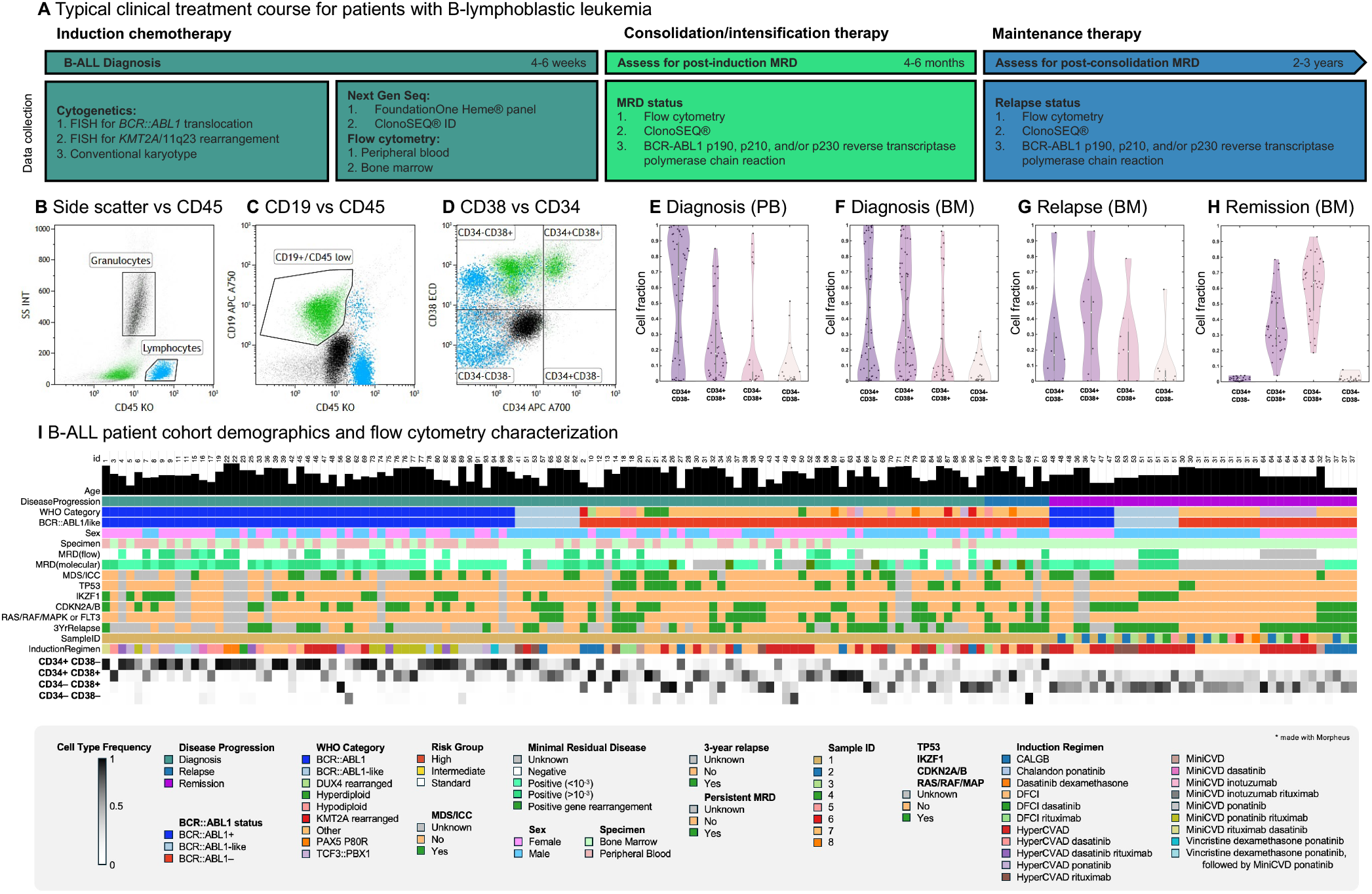
Data collection for B-cell acute lymphoblastic leukemia (B-ALL). **A**. Diagnostic flow cytometry, molecular genetics, and cytogenetic data were obtained for each patient prior to beginning induction chemotherapy and included the elements listed above. Minimal residual disease test results were also obtained throughout each patient’s clinical course, and instances of relapse and death were documented to determine the potential impact of B-ALL subpopulation dynamics on therapy response and clinical outcome. **B.-D**. A simple flow cytometry gating strategy was used to select the leukemia population and divide it into CD34^+^/CD38^-^ (hematopoietic stem cell-like), CD34^+^/CD38^+^ (hematogone-like, stage 1), CD34^-^/CD38^+^ (hematogone-like, stages 2 and 3), and CD34^-^/CD38^-^ (naïve B-cell-like) subpopulations. Neutrophils served as a reference CD34^-^/CD38^-^ population. **E.-H**. The distribution of these subpopulations across the data set is presented using violin plots. **I**. B-ALL patient cohort demographics and flow cytometry characterization at three stages in disease progression: diagnosis, relapse, and remission. Demographic information (sex, age) is shown for patients with matched clinical outcomes (minimal residual disease status and three year relapse), genetic mutations (BCR::ABL1, TP53, IKZF1, CDKN2A/B, RAS/RAF/MAPK pathway or FLT3), WHO Category, sample type, and flow cytometry characterization (CD34+/CD38-, CD34+/CD38+, CD34-/CD38+, CD34-/CD38-). MRD is defined using multiple modalities (molecular/ flow).

The flow cytometry gating strategy that was applied to leukemia cells is shown in **Figure 1B-D**. Mature neutrophils and lymphocytes, identified from side scatter versus CD45, served as reference populations to gauge antigen expression. The leukemia population was selected using a polygonal gate to include all CD19-positive, CD45-dim positive to CD45-negative cellular events. Then, it was divided into quadrants based on CD34 and CD38 expression with neutrophils acting as a reference CD34^-^/CD38^-^ population. In this way, each B-ALL was described as a composite of four cell states, as follows: CD34^+^/CD38^-^ (hematopoietic stem cell-like), CD34^+^/CD38^+^ (hematogone-like, stage 1), CD34^-^/CD38^+^ (hematogone-like, stages 2 and 3), and CD34^-^/CD38^-^ (naïve B-cell-like). **Figure 1E-H** shows these subpopulations in B-ALL patient samples from bone marrow (N=63) and peripheral blood (N=46), as well as bone marrow samples from a cohort of individuals in molecular remission for comparison.

The dataset is summarized in **Table 1** and visualized in **Figure 1I**, where each row contains patient clinical information (e.g. age, genomic markers, specimen type), along with flow cytometry characterization (CD34 and CD38 status). Each column is an individual patient, grouped from left to right into samples taken at diagnosis, relapse, or remission. The median age of the cohort was 55 years old (interquartile range, IQR, 41-70). There was a balanced mix of and *BCR::ABL1* positive versus negative samples and peripheral blood versus bone marrow samples.. All patients had a sample collected at diagnosis, and 32 (34%) had a sample from one or two additional clinical time points. Samples from patients in remission provided a comparator group for the leukemic patients at diagnosis or relapse. At diagnosis, *BCR::ABL1* positive and *BCR::ABL1*-like patients had a higher proportion of CD34^+^CD38^-^ cells, while *BCR::ABL1* negative patients typically had a lower proportion of CD34^+^/CD38^-^ cells and higher CD34^+^/CD38^+^ or CD34^-^/CD38^+^ cell fraction. Patients in remission displayed a high frequency of CD34^-^/CD38^+^ cells.

**Table 1.**
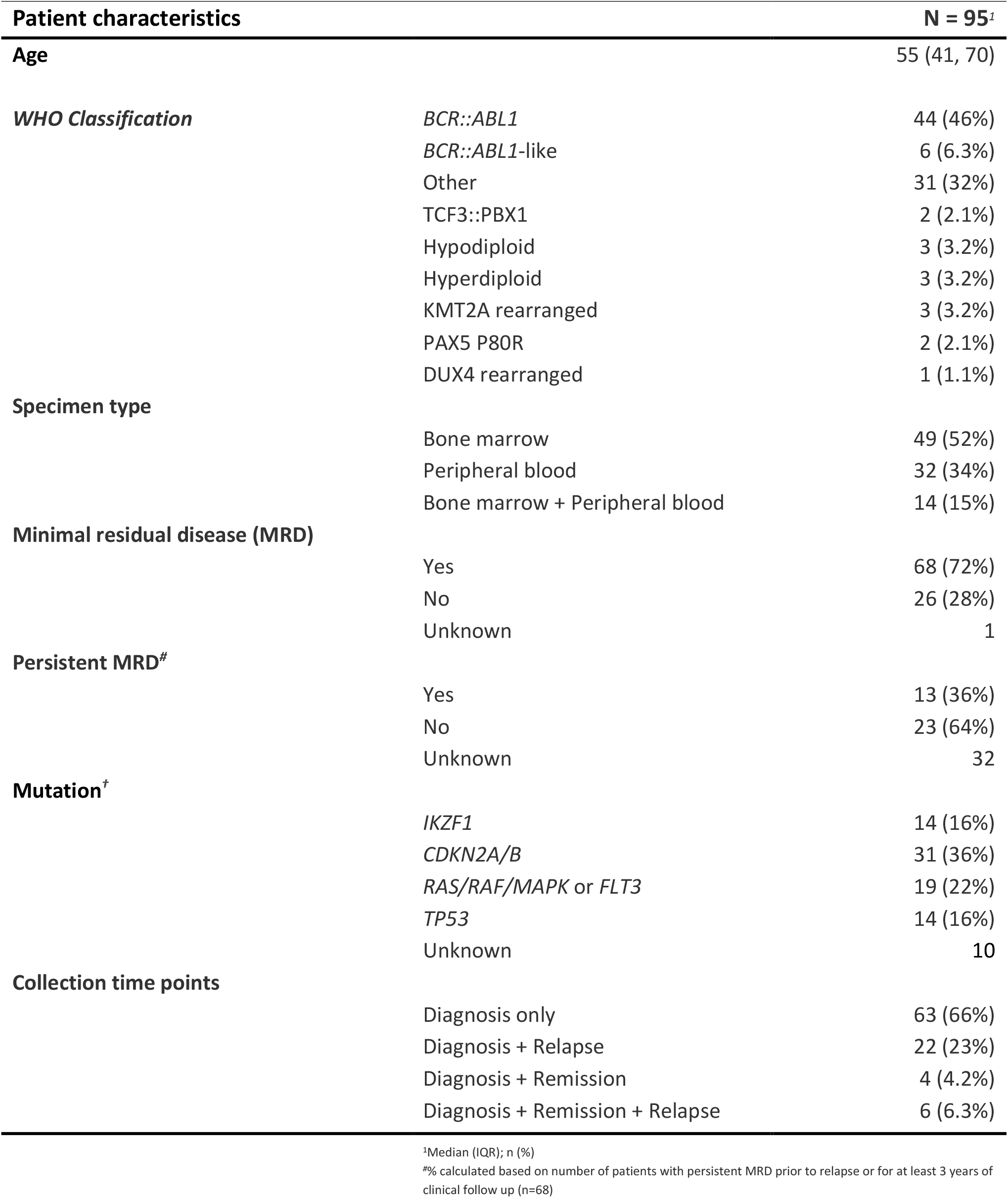
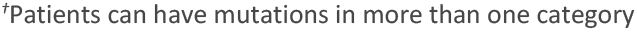
Description of the B-ALL patient cohort.

| Patient characteristics |  | N = 95<sup>1</sup> |
| --- | --- | --- |
| Age |  | 55 (41, 70) |
| **WHO Classification** | *BCR::ABL1* | 44 (46%) |
|  | *BCR::ABL1*-like | 6 (6.3%) |
|  | Other | 31 (32%) |
|  | TCF3::PBX1 | 2 (2.1%) |
|  | Hypodiploid | 3 (3.2%) |
|  | Hyperdiploid | 3 (3.2%) |
|  | KMT2A rearranged | 3 (3.2%) |
|  | PAX5 P80R | 2 (2.1%) |
|  | DUX4 rearranged | 1 (1.1%) |
| **Specimen type** | Bone marrow | 49 (52%) |
|  | Peripheral blood | 32 (34%) |
|  | Bone marrow + Peripheral blood | 14 (15%) |
| **Minimal residual disease (MRD)** | Yes | 68 (72%) |
|  | No | 26 (28%) |
|  | Unknown | 1 |
| **Persistent MRD<sup>#</sup>** | Yes | 13 (36%) |
|  | No | 23 (64%) |
|  | Unknown | 32 |
| **Mutation<sup>†</sup>** | *IKZF1* | 14 (16%) |
|  | *CDKN2A/B* | 31 (36%) |
|  | *RAS/RAF/MAPK* or *FLT3* | 19 (22%) |
|  | *TP53* | 14 (16%) |
|  | Unknown | 10 |
| **Collection time points** | Diagnosis only | 63 (66%) |
|  | Diagnosis + Relapse | 22 (23%) |
|  | Diagnosis + Remission | 4 (4.2%) |
|  | Diagnosis + Remission + Relapse | 6 (6.3%) |

<sup>1</sup>Median (IQR); n (%)
<sup>#</sup>% calculated based on number of patients with persistent MRD prior to relapse or for at least 3 years of clinical follow up (n=68)

### Patients in molecular remission are associated with low CD34^+^CD38^-^ cell self-renewal

We employ a Markov chain-based mathematical modeling approach to estimate cell state transition rates among four distinct cell states, characterized by CD34 and CD38 flow cytometry markers (see **Methods** for details on model parameterization). The model encompasses 16 possible transitions, accounting for all theoretically possible connections between the four cell states (though many transitions may be negligible). Each transition is denoted as M_ij_, where i represents the target state and j represents the initial state. For example, M_11_ corresponds to self-renewal within state 1 (the stem cell-like state).

We begin by estimating cell state transition rates for patients in molecular remission, visualized using a chord diagram in **Figure 2A**, which shows the mean transition matrix or typical cell state kinetics, averaged across all remission patients.

**Figure 2:**
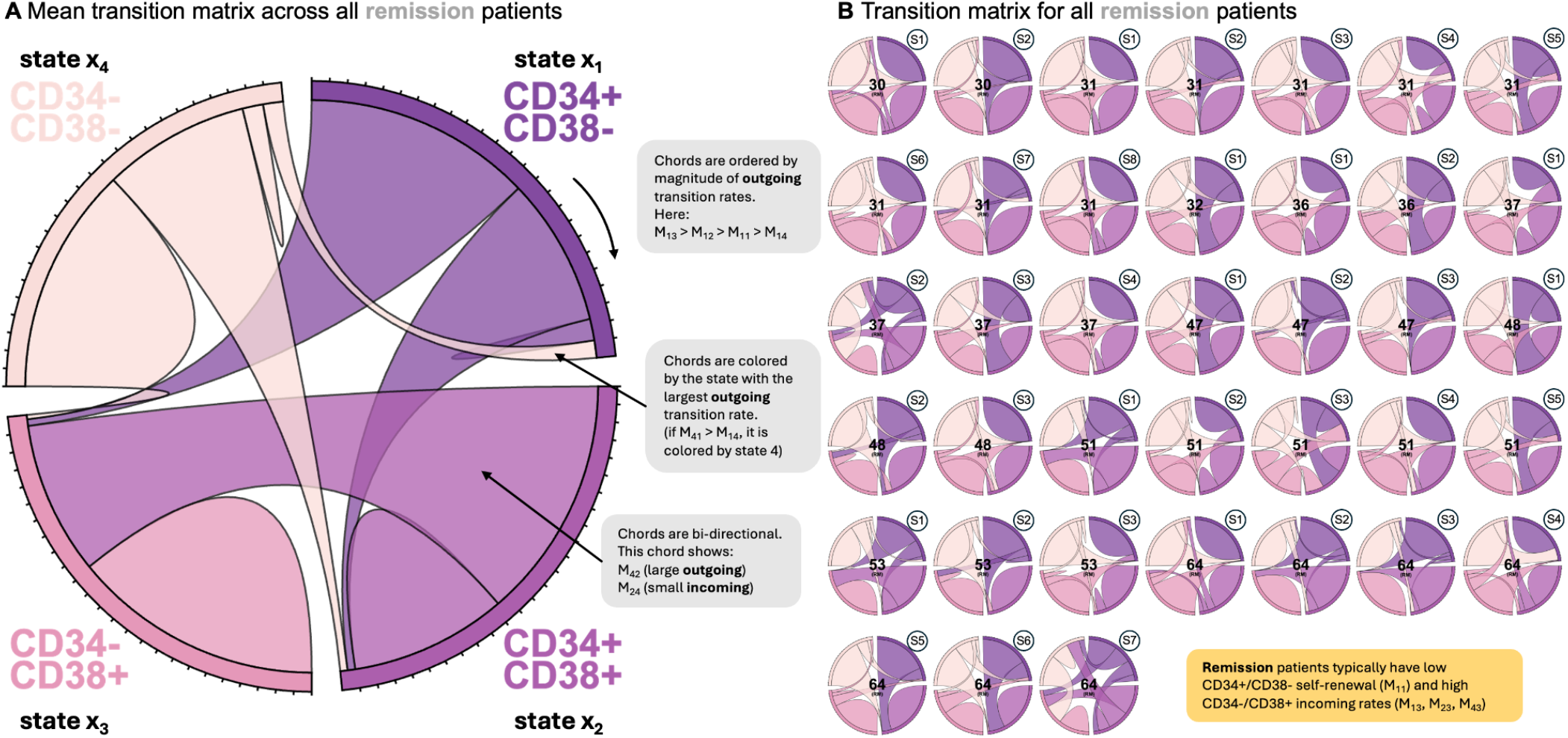
Samples from patients in molecular remission. **A**. Chord diagram illustrating the mean transition matrix across all patients. The chord is organized into four sections (CD34^+^/CD38^-^, CD34^+^/CD38^+^, CD34^-^/CD38^+^, CD34^-^/CD38^-^), with the size of each chord proportional to the magnitude of the outgoing transition rate. Chords are ordered (clockwise) by the magnitude of outgoing transition by each section for visual interpretation, and each bi-directional chord is colored by which is greater: M_ij_ or M_ji_. **B**. Chord diagram shown for each individual patient sample. Sample ID number is shown at the top right of each chord diagram.

As the CSC hypothesis is well supported in AML, we begin by making observations on transitions to and from the hematopoietic stem cell compartment (CD34^+^/CD38^-^) in this B-ALL dataset. Patients in remission typically have a low CD34^+^/CD38^-^ renewal rate, M_11_. This low degree of CD34^+^/CD38^-^ cell self-renewal is generally paired with a high degree of transitions into the CD34^-^ /CD38^+^ cell state and reflects the normal maturation of hematogones in bone marrow. This provides a baseline for the cell state transitions to compare with leukemia patients. We also note that there is a moderate degree of inter-patient heterogeneity across all patients (**Figure 2B**).

### Matched peripheral blood and bone marrow samples

Next, we estimate cell state transitions for samples from B-ALL patients at diagnosis (**Figure 3**). The mean transition matrix (**Figure 3A**) for *BCR::ABL1* positive patients is typically associated with high CD34^+^/CD38^-^ cell self-renewal (M_11_) and high CD34^+^/CD38^-^ incoming rates (M_21_, M_21_, M_31_). In contrast, *BCR::ABL1* negative patients have low CD34^+^/CD38^-^ cell self-renewal rates (M_11_) and one of the following characteristics: high CD34^+^/CD38^+^ incoming rates (M_12_, M_32_, M_42_) or high CD34^-^/CD38^+^ incoming rates (M_13_, M_23_, M_43_). Visually, there is a moderate degree of inter-patient heterogeneity across all samples (**Figure 3B**).

**Figure 3:**
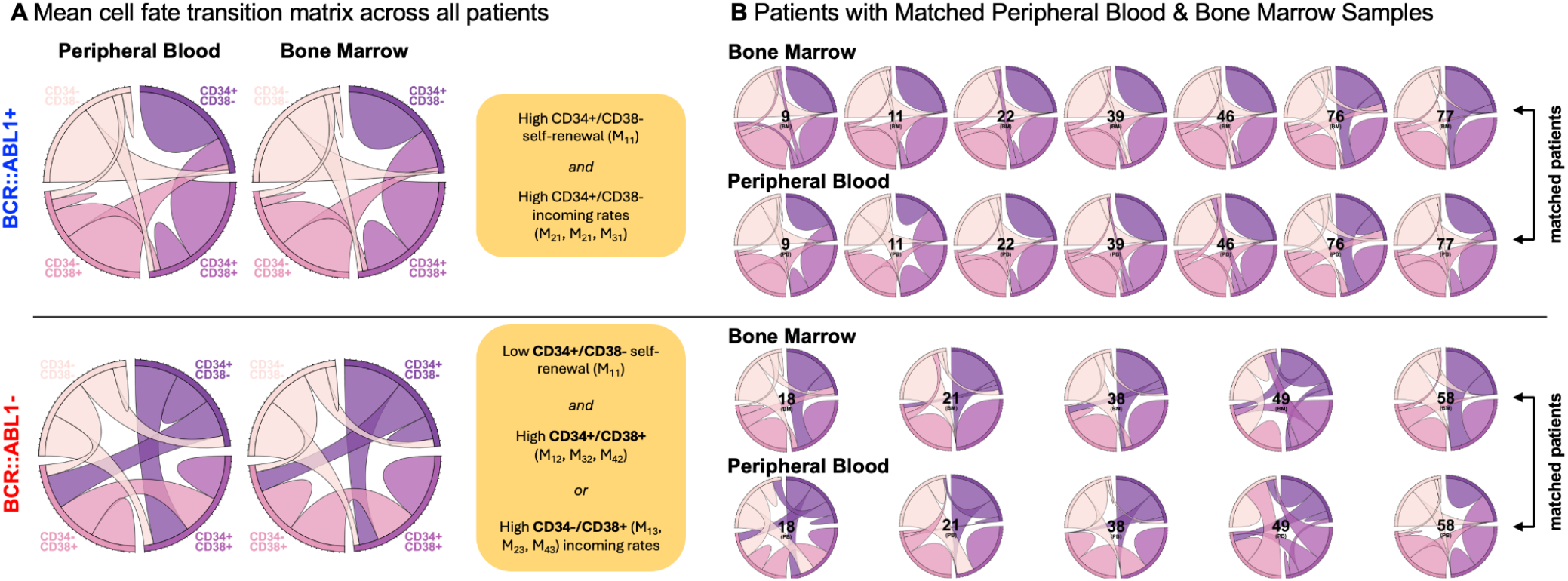
Samples from leukemia patient samples. **A**. Chord diagram illustrating the mean transition matrix across all patients, for all peripheral blood (PB, left) and bone marrow samples (BM, right), repeated for *BCR::ABL1* positive (top) and *BCR::ABL1* negative (bottom) patients. *BCR::ABL1*+ samples are typically associated with high CD34^+^/CD38^-^ self-renewal (M_11_) and high CD34^+^/CD38^-^ incoming rates (M_21_, M_21_, M_31_). *BCR::ABL1* negative patient samples have low CD34^+^/CD38^-^ self-renewal rates (M_11_) and either 1) high CD34^+^/CD38^+^ (M_12_, M_32_, M_42_) or 2) high CD34^-^/CD38^+^ (M_13_, M_23_, M_43_) incoming rates. **B**. Matched samples from PB and BM reach good agreement in transition rates for both *BCR::ABL1* positive or negative samples.

A subset of patients has matched peripheral blood (PB) and bone marrow (BM) leukemia samples, enabling direct comparison of specimen types (**Figure 3B**). Model-predicted cell state transition kinetics are remarkably similar across PB and BM leukemia samples within the same patient, and thus we are unable to reject the null hypothesis that PB and BM samples are drawn from an identical distribution for all 16 transition rate parameters (see **Supplemental Figure S1**). Therefore, for the rest of the analyses we will group PB and BM samples together.

### Stem cell hypothesis revisited

Next, we observe the relationship between important cell state transition values for each patient and correlate salient findings with patient classifications (*BCR::ABL1* status) and clinical outcomes (MRD and relapse). Beginning with *BCR::ABL1*, we use a principal component analysis (PCA) to assess the ability of the 16 Markov transition parameters in clustering patients based on *BCR::ABL1* status (**Figure 4A**). The PCA projects each patient sample onto two axes (PC1, PC2) which are a linear combination of all 16 input features (the 16 Markov transition parameters). The PCA technique visualizes the contribution of each input parameter (M_ij_) as a vector, indicating how that parameter influences each principal component. We can make several important observations from this PCA. The vectors (**Figure 4A**, black arrows) representing transition Matrix parameters cluster tightly with parameters from the same *incoming* compartment. For example, M_11_ clusters with the CD34^+^/CD38^-^ cell state’s other incoming parameters: M_21_, M_31_, and M_41_. This means that if one of these parameters is high (or low), then the other three are likely to be equally high (or low). We note that the PCA provides a reasonable separability between *BCR::ABL1* positive (blue) and *BCR::ABL1* negative patients. Strikingly, remission patients have much lower transition rates into the CD34^+^/CD38^-^ and CD34^-^/CD38^+^ cell states than corresponding samples from leukemia patients not in remission.

**Figure 4:**
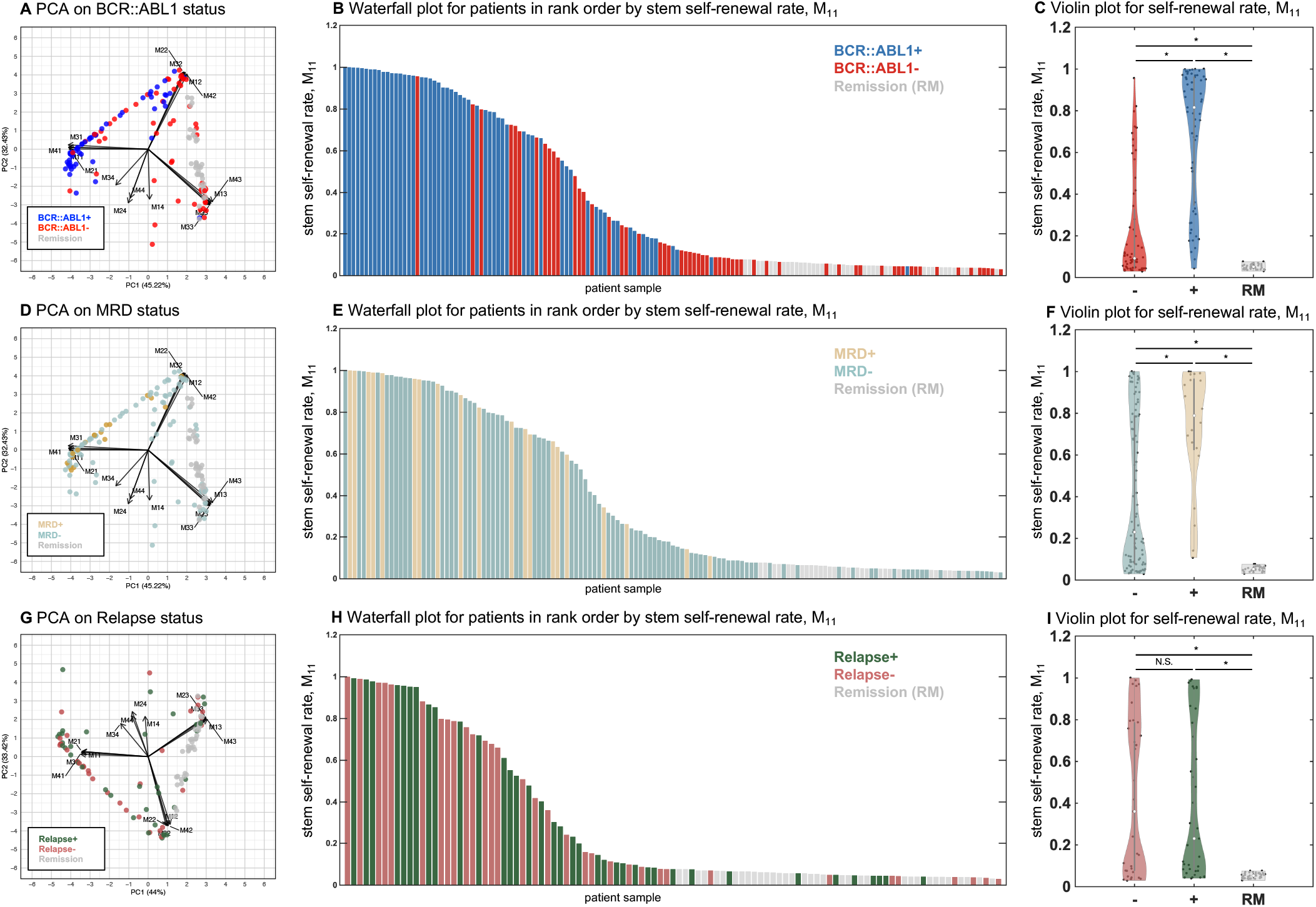
Self-renewal transition rate correlates with *BCR::ABL1* status and Minimal Residual Disease. **A**. Principle Component Analysis (PCA) of all Markov transition rate parameters, M_ij_, color-coded by *BCR::ABL1* status. Arrows indicate the loadings for each parameter, representing the correlation between the variables and the principal components. *BCR::ABL1*+ patients tend to cluster near loadings for transition parameters back into the stem cell state, M_11_, M_21_, M_31_, and M_41_. **B**. Waterfall plot for patients in rank order by value of stem cell self-renewal transition rate, M_11_. **C**. Violin plot for self-renewal rate, M_11_, categorized by *BCR::ABL1* status, showing statistically different values. **D**. PCA color-coded by MRD outcomes. **D**. Waterfall plot rank ordered by stem self-renewal transition rate, M_11_, color-coded by MRD outcome. **F**. Violin plot for stem self-renewal transition rate, M_11_, color-coded by MRD outcome. **G**. PCA color-coded by Relapse. **H**. Waterfall plot rank ordered by stem cell self-renewal transition rate, M_11_, color-coded by Relapse. **I**. Violin plot for stem cell self-renewal transition rate, M_11_, color-coded by Relapse.

Because all incoming cell state transition parameters tend to cluster together, we need only to observe the self-transition parameter M_11_, to gain insight into transitions toward the CD34^+^/CD38^-^ cell state. The waterfall plot shown in **Figure 4B** shows a consistent rank-order relationship of *BCR::ABL1* positive (high M_11_), *BCR::ABL1* negative (mid-range M_11_), to remission samples (low M_11_). The violin plot confirms the statistical significance of the relationship between self-renewal rate and *BCR::ABL1* status, with a baseline of comparison to molecular remission patients also shown. Violin plots for all rates are shown in **Supplemental Figure S2**.

Next, we repeat the PCA and waterfall for post-induction chemotherapy MRD status (**Figure 4D-F**) and three-year disease relapse status (**Figure 4G-I**). Here, MRD is defined by a threshold frequency of 10-3 leukemia cells detected using specialized flow cytometry and relapse is defined as morphologic relapse. A high self-renewal transition rate also correlates to an increased likelihood of positive MRD (**Figure 4F**). However, the relationship between cell state transitions and relapse does not show any significant correlations (**Figure 4I**), consistent with previous literature attempting to predict relapse/recurrence in childhood B-ALL^47,48^.

### Matched diagnosis and relapse samples indicate relatively stable cell state kinetics

We hypothesized that the lack of correlation between cell state transitions and relapse may be due to time-varying transition values. We sought to understand if disease cell state kinetics remain stable before and after treatment for patients who relapse. To address this, we also trained the model using patient samples taken at the point of relapse. This allows us to directly compare cell state transition rates between diagnosis and relapse time points. Although there is significant variation among patients (**Figure 5A**), we do not observe any clear, statistically significant patterns that correlate between diagnosis and relapse (**Figure 5B**). In contrast, matched samples from patients in molecular remission (**Figure 5C**) consistently show clear differences in cell state transition rates (**Figure 5D**), particularly in transitions back to compartments 1 (CD34^+^/CD38^-^) and 3 (CD34^-^/CD38^+^). Incoming CD34^+^/CD38-transition rates are lower in remission patients (compared to diagnosis baseline) while incoming CD34^-^/CD38^+^ transition rates are higher value in remission patients.

**Figure 5:**
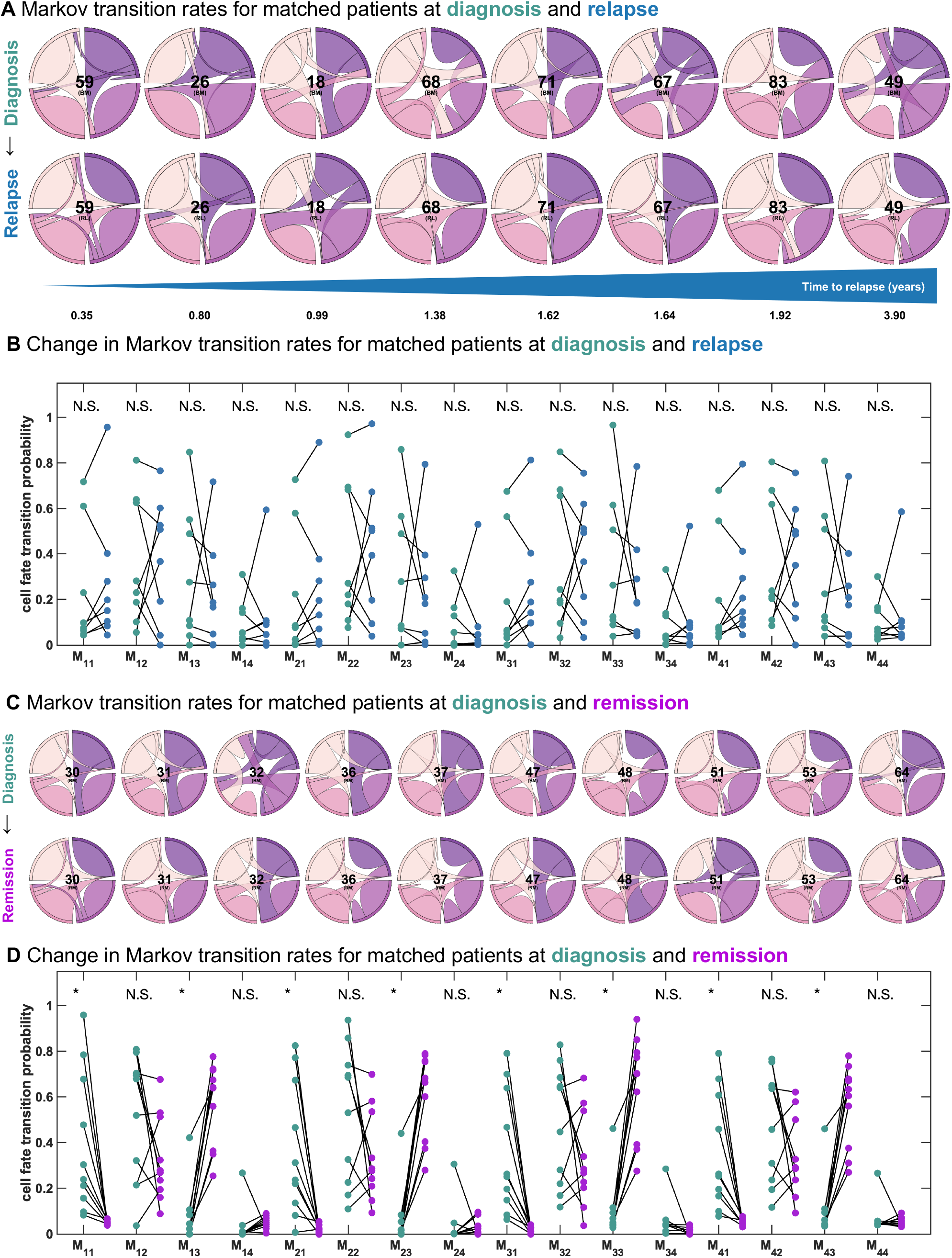
Matched samples from patients at diagnosis and relapse. **A**. Markov transition rate chord diagrams for matched patients at diagnosis (top) and relapse (bottom), ordered from left to right by time to relapse. **B**. Biplot showing the change in each transition parameter value at diagnosis and relapse (left to right). Note all 16 transition parameters are not significantly different, indicating no clear pattern of change between diagnosis and relapse across our cohort of matched samples **C**. Markov transition rate chord diagrams for matched patients at diagnosis (top) and remission (bottom). **D**. Biplot showing the change in each transition parameter value at diagnosis and remission (left to right). Significant changes are observed in all transitions back into compartments 1 and 3 (CD34^+^/CD38^-^ and CD34^-^/CD38^+^, respectively).

We remind the reader that these remission patients are distinctly separable from diagnosis samples, as shown in **Figure 4A, D**, and **G** and **Figure 5D**. Taken together, this suggests that if particular cell state transitions are altered due to therapy, then therapy is more likely to be successful. Remission patients have statistically different values for transitions into compartments 1 (CD34^+^/CD38^-^) and 3 (CD34^-^/CD38^+^), but this change does not occur for relapse-matched patients. We note that the question of whether or not transition rates change between diagnosis and relapse (**Figure 5**) is a different question than whether the rates at the point of diagnosis predict which patients are more likely to relapse (**Figure 4G-I**).

## Discussion

In this work, we employed a Markov chain mathematical model to quantify the rates of cell state transitions between CD34^+^/CD38^-^, CD34^+^/CD38^+^, CD34^-^/CD38^+^, and CD34^-^/CD38^-^ leukemia subpopulations in patients with B-ALL. Such transitions have been observed in vitro using time-lapse fluorescence microscopy and cell sorting but have not been captured in the clinical laboratory, primarily because diagnostic flow cytometry and immunohistochemistry represent a mere snapshot of the malignant population at a single point in time. Our method can infer subpopulation dynamics for any hematolymphoid malignancy where clinical flow cytometry data is available. It may also be possible to apply a similar approach wherever quantitative markers are reported in pathology; for instance, hormone receptor expression in breast carcinoma or PD-L1 expression in non-small cell lung carcinoma.

In B-ALL, we find that subpopulation dynamics are useful to predict *BCR::ABL1* status and post-induction MRD (threshold frequency of 10^−3^ leukemia cells detected using specialized flow cytometry). Our analysis indicates the existence of de-differentiation transitions to a CD34^+^/CD38^-^ stem cell-like immunophenotype in leukemia samples collected from B-ALL patients, and the tendency for these transitions is especially strong for B-ALL with *BCR::ABL1*. Also of note, transition values were found to be non-zero for all but one patient (see **Supplemental Tables T1-T3**). We interpret these findings as contrary to the hierarchical nature of the cancer stem cell hypothesis. Instead, these results favor a non-hierarchical relationship between the leukemia stem cell compartment (CD34^+^/CD38^-^) and other cell states, such that it is possible to access all cell states from any cell state. This has implications for therapies targeting cancer stem cells because CSCs would be replenished over time by transitions from non-stem cells unless the transitions are somehow inhibited.

The results of the study has practical uses for pathologists and oncologists at present, such as predicting *BCR::ABL1* when a sample is inadequate for molecular genetic and cytogenetic studies. Since *BCR::ABL1*-like patients cluster with *BCR::ABL1*-positive patients in our analysis, it may also be possible to predict *BCR::ABL1*-like acute leukemias without testing for specific kinase-activating rearrangements such *IGH*::*CRLF2*, as long as the *BCR::ABL1* status is known. Other genotypic findings of less clinical significance also show significant associations with the model results. It is known that *BCR::ABL1* and *TP53* mutations are almost mutually exclusive in acute lymphoblastic leukemias and, interestingly, patients with these two genetic findings in our study generally have distinct immunophenotypes: a dominant CD34^+^/CD38^-^ subpopulation if *BCR::ABL1*-positive, versus a combination of CD34^+^/CD38^+^ and CD34^-^/CD38^+^ cells if *TP53* mutant.

Importantly, the model parameters were unable to predict relapse after three years. We note that this null result is consistent with other findings in literature^47^, and suggests that temporal disease evolution occurring during the treatment course may not be reflected in routine clinical flow cytometry data. Our original hypothesis was that cell states (CD34^+^/CD38^-^, CD34^+^/CD38^+^, CD34^-^/CD38^+^, and CD34^-^/CD38^-^) may have different survival rates during therapy and so we should observe selection toward a subpopulation equilibrium that maximizes transitions to the most therapy-resistant cell state, while minimizing transitions to the least resistant cell state. There were no statistically significant differences between cell state transitions in paired diagnosis and relapse specimens, but it is unclear whether this is due to the relatively small number of eligible paired specimens we were able to identify, or perhaps that the subpopulations do not differ significantly in terms of fitness under treatment-based selection. Although leukemia stem cells have been reported to increase in frequency during treatment in AML^8^, CD34 and CD38 were not among the markers observed to increase, raising a further possibility that our choice of markers precluded the observation of treatment-based selection. The application of our method to additional markers and types of neoplasia may yet produce clinically significant results.

The primary achievement of the study is the successful implementation of a Markov model to infer cell state transitions, which would otherwise go unstudied due to the cross-sectional nature of immunophenotyping in clinical laboratories. Our modeling approach also suggests that targeting CD34^+^/CD38^-^ self-renewal rate (i.e. reducing it) would represent a promising treatment strategy. Patients in molecular remission consistently exhibit lower CD34^+^/CD38^-^ and CD34^-^ /CD38^+^ values compared to diagnosis. Notably, these transition rates differ between patients who achieve remission and those who relapse. This observation suggests that transition rates change due to the course of therapy, and we hypothesize that both the CD34^+^/CD38^-^ and CD34^-^ /CD38^+^ de-differentiation transitions represent a potential therapeutic target to improve remission rates. This is confirmed via statistically significant differences between paired samples from the same patients at both diagnosis and remission time points across all cell state transitions into CD34^+^/CD38^-^ and CD34^-^/CD38^+^. Incoming CD34+/CD38-show a decreasing transition rate in remission patients (compared to diagnosis baseline) while incoming CD34^-^ /CD38^+^ show an increasing value in remission patients. The differences observed between the diagnosis, relapse, and remission groups highlight the critical role of cell state transition rates in patient outcomes.

## Materials & Methods

### Patient inclusion criteria

Following institutional review, patients eligible for inclusion in the study were identified by searching the laboratory information system at Moffitt Cancer Center for all instances of a specific 10-color flow cytometry panel performed between May 2016 and May 2023. Each diagnostic flow cytometry result was reviewed to confirm the relevant panel had been performed on a treatment-naive specimen and the diagnosis was indeed B-ALL in conjunction with clinical and other pathological findings. Then, patients selected for inclusion in the study underwent chart review to obtain information including age, sex, date of diagnosis, molecular genetic, and cytogenetic findings, treatment regimen, MRD test results, date of last contact, and date of relapse and/or death, if applicable.

Genetic data consisted of a hybrid capture-based next-generation sequencing (NGS) panel (FoundationOne Heme), conventional karyotyping, *BCR/ABL1/ASS1* tri-color fluorescence in situ hybridization (FISH), and *MLL* (*KMT2A*) 11q23.3 break-apart FISH as applicable per patient. MRD testing consisted of 10-color flow cytometry, quantitative real-time reverse transcriptase polymerase chain reaction (PCR) for *BCR::ABL1* p190 and p210 fusion transcripts, and multiplex PCR and NGS to identify clonal immunoglobulin gene sequences (ClonoSEQ) or gene fusions as applicable per patient. The 10-color flow cytometry panel used for diagnosis and MRD detection was performed on a Gallios System and analyzed using Kaluza software (Beckman Coulter, IN). Antibodies included in the panel are CD15, CD130, CD10, CD58, CD22, CD33, CD13, CD123, CD19, CD20, CD45, CD38, CD34, CD81, CD200 (BD, Biolegend, Beckman Coulter). A minimum of 750,000 events were collected for MRD assays with a validated lower limit of detection of 0.01%.

### Sample collection and analysis

The proportion of CD34^+^/CD38^-^, CD34^+^/CD38^+^, CD34^-^/CD38^+^, and CD34^-^/CD38^-^ leukemia cells in diagnostic peripheral blood and/or bone marrow specimens was determined using a simple flow cytometry gating strategy that utilized mature neutrophils and lymphocytes as reference populations (**Figure 1**). Samples were first analyzed to exclude doublets and debris. Then, mature neutrophils and lymphocytes were identified from a side scatter versus CD45 plot and painted separate colors. The leukemia population was selected by drawing a polygonal gate to include all CD19-positive, CD45-dim positive to negative events and, if necessary, plasma cells were excluded using an interdependent gate to remove CD38-bright events. Lastly, all cellular events were visualized on a CD34 versus CD38 plot, and the leukemia population was divided into CD34^+^/CD38^-^, CD34^+^/CD38^+^, CD34^-^/CD38^+^, and CD34^-^/CD38^-^ quadrants with mature neutrophils serving a CD34^-^/CD38^-^ reference population.

### Markov chain model parameterization

We next developed a Markov chain mathematical model to quantify transition rates between the four leukemia cell states defined by CD34 and CD38 expression (**Figure 6**). We employed the Newton-Mason iterative, algorithmic search process^49,50^, using flow cytometry characterization of four B-ALL cell states stated above. This numerical search procedure was used to derive patient-specific Markov matrices, describing the stochastic cell state transitions and resulting in estimates for cell state transition rates for each patient. A numerical Markov chain modeling training procedure was performed on flow cytometry data of a cohort of B-ALL patient samples of peripheral blood (N=46) and bone marrow (N=63) with matched clinical and molecular features such as *BCR::ABL1* status, comprehensive genomic profiling, minimal residual disease (MRD) post-induction chemotherapy, and 3-year relapse. Critical to our goal of quantifying the evolution of state transition rates, we also obtained bone marrow measurements for a cohort of individuals in molecular remission (N=38).

**Figure 6:**
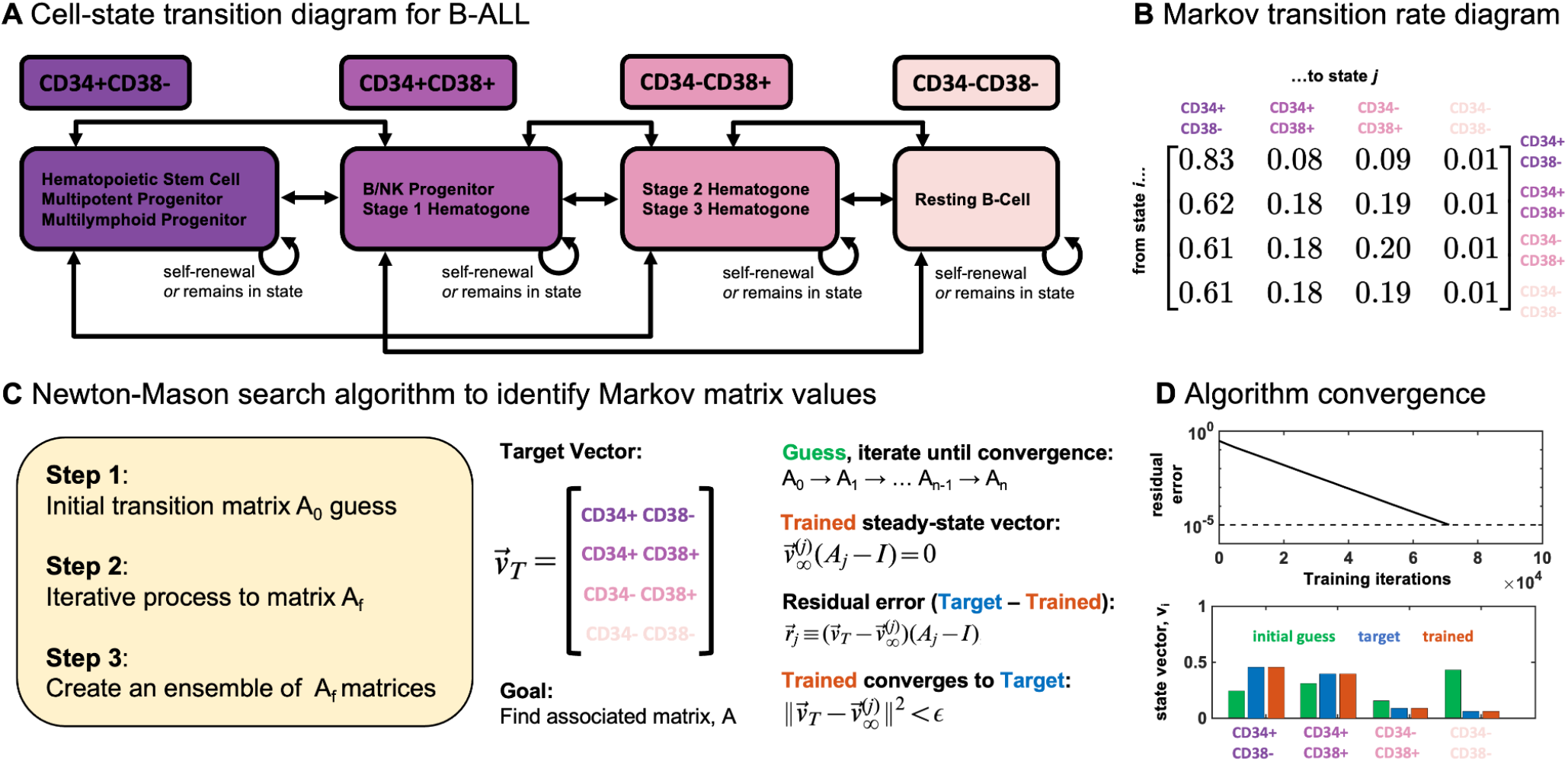
Parameterization process for Markov chain model. **A**. Schematic of cell states characterized by CD34 and CD38, and the 16 possible transitions between 4 states. **B**. Markov transition rate matrix which represents the transition probability from the state given by the row, into the state given by the column. **C**. Newton-Mason search algorithm identifies Markov matrix parameter values in three steps: initial guess, iteration, and ensemble. **D**. Validation of algorithm convergence. Initial guess has a high residual error, which converges logarithmically toward the target vector until acceptable tolerance is reached (dashed line).

### Markov transition matrix identification

We describe the transition of cells between states using a stochastic transition matrix, *M*, within a Markov dynamical system. The proportion of cells in each of *i* states is contained in the state-vector 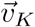 for discrete time steps, *K*, such that the state changes according to the following equation:

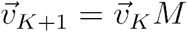

A well-known property of Markov dynamic systems is that the state-vector at time step *K* can be found directly by:

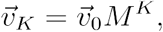

where 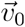 is the initial state-vector at time zero. Each matrix has a corresponding steady-state vector defined as:

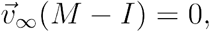

where *I* is the identity matrix. Here, our state vector contains 4 states where index *i* ∈ [1, 2, 3, 4] corresponds to CD34^+^/CD38^-^, CD34^+^/CD38^+^, CD34^-^/CD38^+^, CD34^-^/CD38^-^, respectively (see **Figure 6A,B**).

### Algorithm to compute Markov transition matrix

Next, we use an iterative search algorithm (**Figure 6C**) to determine the most likely Markov matrix, *M*, associated with each patient-specific target vector, 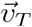. This target vector composition is determined by flow cytometry for each patient at diagnosis before induction chemotherapy. We assume that the target vector represents quasi-steady state disease dynamics, and thus the search algorithm finds a Markov matrix such that the follow relation is true:

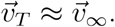

Following the process outlined by Newton et. al.^49,50^, there is a three-step process to compute the transition matrix, *M*, which we restate below.

**Step 1**: the initial choice of Markov matrix, *M*. The Newton-Mason training algorithm is non-ergodic, and thus depends on the initial condition chosen to begin the iterative search. As previously outlined by Newton et. al.^49,50^, the initial choice should be a rank 2 matrix, where the row indicating the disseminating phenotype (in our case, the stem-like cell type 1) should match the steady-state distribution of disease samples. Stated formally:

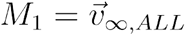

where
is the *M* 1st row of M and 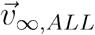 is the steady-state average associated with ALL patients. For all other rows *j* ≠ 1:

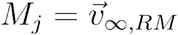

Where 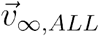 is the steady-state average associated with molecular remission patients.

The intuition behind this choice is that we begin with a matrix where stem cells are assumed to transition to other cells at a rate proportional to each cell type in ALL patients, and non-stem cells transition at rate proportional to each cell type in disease-free molecular remission patients.

**Step 2**: the iteration process to final Markov matrix, *M*_*f*_.Starting with initial Markov matrix *M*_0_, entries are iteratively adjusted by randomized adjustments until the steady-state vector converges to the patient’s target distribution, within some error tolerance. First, the residual error defined at each step *j* of the training process is the difference between the patient’s target vector and the steady-state:

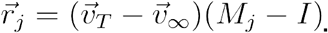

This residual error vector determines the subsequent adjustments made to the Markov matrix for algorithm step *j+1*. Select a random row of M to modulate. A value of *δ* is <u>subtracted</u> from the column of M corresponding to the <u>maximum</u> entry of 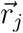, and the same value of *δ* is <u>added</u> from the column of M corresponding to the <u>minimum</u> entry of 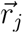. The added and subtracted value *δ* is scaled with the size of the Euclidean norm of the residual vector:

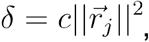

where *c* is a small constant (here, *c* = 0.0005).The search algorithm iterates *M*_*j*_ until numerical convergence threshold such that: 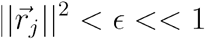, where typically we set *ϵ* = 10^−5^ (**Figure 6D**).

**Step 3**: creating an ensemble of *M*_*f*_ matrices. The algorithm presented in step 2 is randomized (due to randomized row selection), and thus the training process is inherently stochastic. As reported previously, the entries of *M*_*f*_ should be thought of as having an associated probability distribution, with a sample mean and variance obtained by training an ensemble of matrices. We repeat the training process, with identical *M*_0_ many times (typically *N* = 100) and average the Markov entries to obtain the sample means.

## Acknowledgements

The authors gratefully acknowledge funding by the GME Research Grant via the University of South Florida and Moffitt Cancer Center support from the Center of Excellence for Evolutionary Therapy.

## Supplemental Information

**Supplemental Figure S1:**
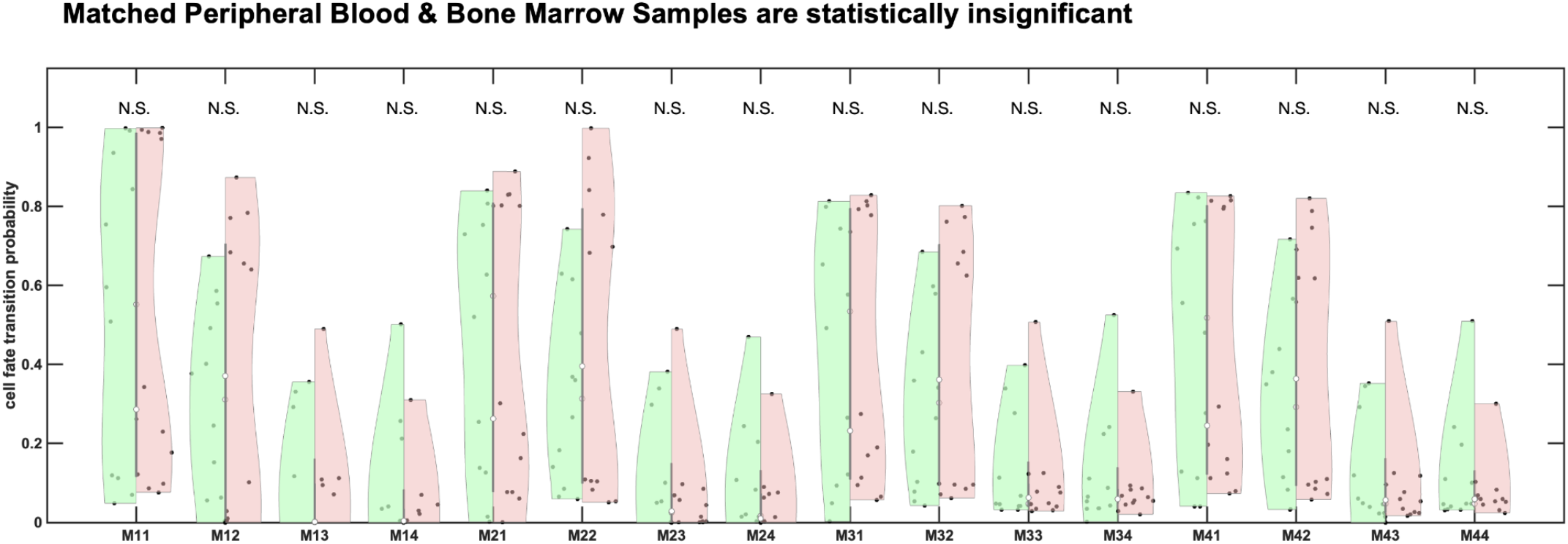
Matched samples from Bone Marrow (green) and Peripheral Blood (red), testing the null hypothesis that the two specimen types come from independent random samples from normal distributions with equal means and equal but unknown variances (two-sample t-test) in our dataset. The null hypothesis is not rejected for all 16 parameters, and marked as not significantly different (N.S) for each.

**Supplemental Figure S2:**
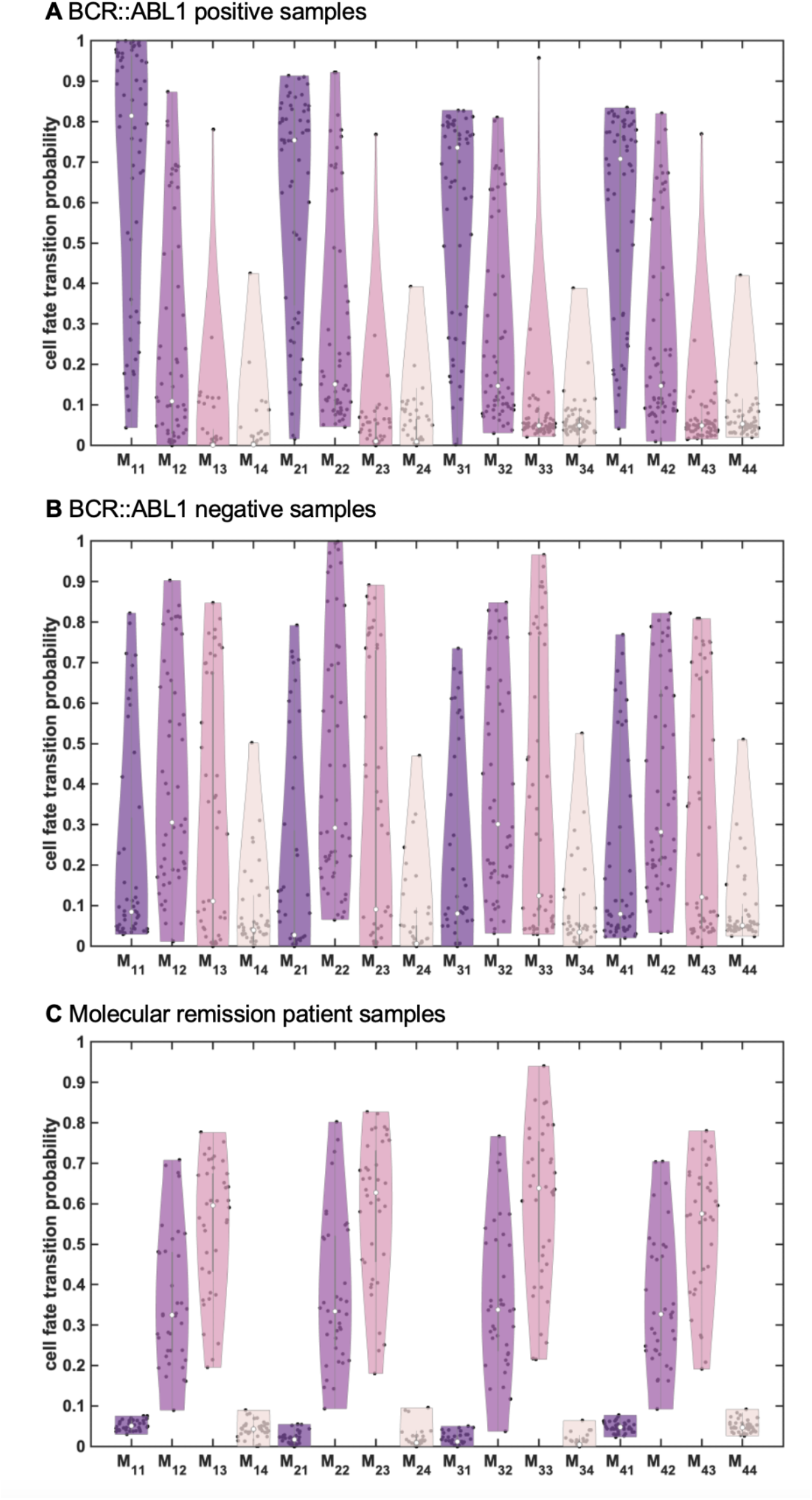
Violin plots of Markov cell state transition rate parameters for (A) BCR::ABL1 positive patient samples, (B) BCR::ABL1 negative patient samples, and (C) molecular remission patient samples.

**Supplemental Table T1:** Table shows the number of remission patient samples that have the corresponding Markov transition parameter statistically significantly equivalent to zero (one-sided t-test).

| Remission<br>(N=38) | Fraction of samples with zero transition rates |  |  |  |
| --- | --- | --- | --- | --- |
|  | CD34<sup>+</sup>/CD38<sup>-</sup> | CD34<sup>+</sup>/CD38<sup>+</sup> | CD34<sup>-</sup>/CD38<sup>+</sup> | CD34<sup>-</sup>/CD38<sup>-</sup> |
| CD34<sup>+</sup>/CD38<sup>-</sup> | 0 | 0 | 0 | 1 |
| CD34<sup>+</sup>/CD38<sup>+</sup> | 2 | 0 | 0 | 6 |
| CD34<sup>-</sup>/CD38<sup>+</sup> | 2 | 0 | 0 | 5 |
| CD34<sup>-</sup>/CD38<sup>-</sup> | 0 | 0 | 0 | 0 |

**Supplemental Table T2:** Table shows the number of bone marrow patient samples that have the corresponding Markov transition parameter statistically significantly equivalent to zero (one-sided t-test).

| Bone Marrow<br>(N=63) | Fraction of patients with zero transition rates |  |  |  |
| --- | --- | --- | --- | --- |
|  | CD34<sup>+</sup>/CD38<sup>-</sup> | CD34<sup>+</sup>/CD38<sup>+</sup> | CD34<sup>-</sup>/CD38<sup>+</sup> | CD34<sup>-</sup>/CD38<sup>-</sup> |
| CD34<sup>+</sup>/CD38<sup>-</sup> | 0 | 3 | 10 | 9 |
| CD34<sup>+</sup>/CD38<sup>+</sup> | 1 | 0 | 2 | 7 |
| CD34<sup>-</sup>/CD38<sup>+</sup> | 2 | 0 | 0 | 3 |
| CD34<sup>-</sup>/CD38<sup>-</sup> | 0 | 0 | 0 | 0 |

**Supplemental Table T3:** Table shows the number of peripheral blood patient samples that have the corresponding Markov transition parameter statistically significantly equivalent to zero (one-sided t-test).

| Peripheral Blood<br>(N=46) | Fraction of patients with zero transition rates |  |  |  |
| --- | --- | --- | --- | --- |
|  | CD34<sup>+</sup>/CD38<sup>-</sup> | CD34<sup>+</sup>/CD38<sup>+</sup> | CD34<sup>-</sup>/CD38<sup>+</sup> | CD34<sup>-</sup>/CD38<sup>-</sup> |
| CD34<sup>+</sup>/CD38<sup>-</sup> | 0 | 1 | 11 | 7 |
| CD34<sup>+</sup>/CD38<sup>+</sup> | 0 | 0 | 3 | 2 |
| CD34<sup>-</sup>/CD38<sup>+</sup> | 0 | 0 | 0 | 0 |
| CD34<sup>-</sup>/CD38<sup>-</sup> | 0 | 0 | 1 | 0 |

## References

1. Lapidot T, Sirard C, Vormoor J, et al. A cell initiating human acute myeloid leukaemia after transplantation into SCID mice. Nature. 1994;367(6464):645–648.

2. Bonnet D, Dick JE. Human acute myeloid leukemia is organized as a hierarchy that originates from a primitive hematopoietic cell. Nat. Med. 1997;3(7):730–737.

3. Grouw EPLM de, Raaijmakers MHGP, Boezeman JB, et al. Preferential expression of a high number of ATP binding cassette transporters in both normal and leukemic CD34+CD38 − cells. Leukemia. 2006;20(4):750–754.

4. Stevens BM, Jones CL, Pollyea DA, et al. Fatty acid metabolism underlies venetoclax resistance in acute myeloid leukemia stem cells. Nat. Cancer. 2020;1(12):1176–1187.

5. Terwijn M, Zeijlemaker W, Kelder A, et al. Leukemic Stem Cell Frequency: A Strong Biomarker for Clinical Outcome in Acute Myeloid Leukemia. PLoS ONE. 2014;9(9):e107587.

6. Hanekamp D, Denys B, Kaspers GJL, et al. Leukaemic stem cell load at diagnosis predicts the development of relapse in young acute myeloid leukaemia patients. Br. J. Haematol. 2018;183(3):512– 516.

7. Rhenen A van, Feller N, Kelder A, et al. High Stem Cell Frequency in Acute Myeloid Leukemia at Diagnosis Predicts High Minimal Residual Disease and Poor Survival. Clin. Cancer Res. 2005;11(18):6520–6527.

8. Ho T-C, LaMere M, Stevens BM, et al. Evolution of acute myelogenous leukemia stem cell properties after treatment and progression. Blood. 2016;128(13):1671–1678.

9. Kong Y, Yoshida S, Saito Y, et al. CD34+CD38+CD19+ as well as CD34+CD38−CD19+ cells are leukemia-initiating cells with self-renewal capacity in human B-precursor ALL. Leukemia. 2008;22(6):1207–1213.

10. Jiang Z, Deng M, Wei X, et al. Heterogeneity of CD34 and CD38 expression in acute B lymphoblastic leukemia cells is reversible and not hierarchically organized. J. Hematol. Oncol. 2016;9(1):94.

11. Aoki Y, Watanabe T, Saito Y, et al. Identification of CD34+ and CD34− leukemia-initiating cells in MLL-rearranged human acute lymphoblastic leukemia. Blood. 2015;125(6):967–980.

12. Lazarus HM, Andersen J, Chen MG, et al. Recombinant granulocyte-macrophage colony-stimulating factor after autologous bone marrow transplantation for relapsed non-Hodgkin’s lymphoma: blood and bone marrow progenitor growth studies. A phase II Eastern Cooperative Oncology Group Trial. Blood. 1991;78(3):830–7.

13. Lang F, Wojcik B, Bothur S, et al. Plastic CD34 and CD38 expression in adult B–cell precursor acute lymphoblastic leukemia explains ambiguity of leukemia-initiating stem cell populations. Leukemia. 2017;31(3):731–734.

14. Long J, Liu S, Li K, et al. High proportion of CD34+/CD38−cells is positively correlated with poor prognosis in newly diagnosed childhood acute lymphoblastic leukemia. Leuk. Lymphoma. 2014;55(3):611–617.

15. Ebinger M, Witte K-E, Ahlers J, et al. High frequency of immature cells at diagnosis predicts high minimal residual disease level in childhood acute lymphoblastic leukemia. Leuk. Res. 2010;34(9):1139– 1142.

16. Shman TV, Movchan LV, Aleinikova OV. Frequencies of immature CD34 + CD38 − and CD34 + CD38−CD19 + blasts correlate with minimal residual disease level in pediatric B-cell precursor acute lymphoblastic leukemia. Leuk. Lymphoma. 2013;54(11):2560–2562.

17. Tabernero M, Bortoluci A, Alaejos I, et al. Adult precursor B-ALL with BCR/ABL gene rearrangements displays a unique immunophenotype based on the pattern of CD10, CD34, CD13 and CD38 expression. Leukemia. 2001;15(3):406–414.

18. Watcham S, Kucinski I, Gottgens B. New insights into hematopoietic differentiation landscapes from single-cell RNA sequencing. Blood. 2019;133(13):1415–1426.

19. Sha Y, Wang S, Zhou P, Nie Q. Inference and multiscale model of epithelial-to-mesenchymal transition via single-cell transcriptomic data. Nucleic Acids Res. 2020;48(17):gkaa725..

20. Buder T, Deutsch A, Seifert M, Voss-Böhme A. CellTrans: An R Package to Quantify Stochastic Cell State Transitions. Bioinform. Biol. Insights. 2017;11:1177932217712241.

21. Gupta PB, Fillmore CM, Jiang G, et al. Stochastic State Transitions Give Rise to Phenotypic Equilibrium in Populations of Cancer Cells. Cell. 2011;146(4):633–644.

22. Strobl M, Gallaher J, Robertson-Tessi M, West J, Anderson A. Treatment of evolving cancers will require dynamic decision support. Annals of Oncology. 2023;

23. West J, Robertson-Tessi M, Anderson ARA. Agent-based methods facilitate integrative science in cancer. Trends in Cell Biology. 2023;33(4):300–311.

24. Bull JA, Byrne HM. The Hallmarks of Mathematical Oncology. P Ieee. 2022;110(5):523–540.

25. Angelini E, Wang Y, Zhou JX, Qian H, Huang S. A model for the intrinsic limit of cancer therapy: Duality of treatment-induced cell death and treatment-induced stemness. PLOS Computational Biology. 2022;18(7):e1010319.

26. Foo J, Basanta D, Rockne RC, et al. Roadmap on plasticity and epigenetics in cancer. Physical Biology. 2022;19(3):031501.

27. Gunnarsson EB, D. S, Leder K, Foo J. Understanding the role of phenotypic switching in cancer drug resistance. Journal of Theoretical Biology. 2020;490:110162.

28. Dénes A, Marzban S, Röst G. Global analysis of a cancer model with drug resistance due to Lamarckian induction and microvesicle transfer. Journal of Theoretical Biology. 2021;527:110812.

29. Zhou JX, Pisco AO, Qian H, Huang S. Nonequilibrium Population Dynamics of Phenotype Conversion of Cancer Cells. PLoS ONE. 2014;9(12):e110714.

30. Pedersen RK, Andersen M, Stiehl T, Ottesen JT. Understanding Hematopoietic Stem Cell Dynamics—Insights from Mathematical Modelling. Curr. Stem Cell Rep. 2023;9(1):9–16.

31. Watson CJ, Papula AL, Poon GYP, et al. The evolutionary dynamics and fitness landscape of clonal hematopoiesis. Science. 2020;367(6485):1449–1454.

32. Moeller ME, Père NVM, Werner B, Huang W. Measures of genetic diversification in somatic tissues at bulk and single-cell resolution. eLife. 2024;12:RP89780.

33. Pedersen RK, Andersen M, Stiehl T, Ottesen JT. Mathematical modelling of the hematopoietic stem cell-niche system: Clonal dominance based on stem cell fitness. Journal of Theoretical Biology. 2021;518:110620.

34. Boklund TI, Snyder J, Gudmand-Hoeyer J, et al. Mathematical modelling of stem and progenitor cell dynamics during ruxolitinib treatment of patients with myeloproliferative neoplasms. Front. Immunol. 2024;15:1384509.

35. Kreger J, Mooney JA, Shibata D, MacLean AL. Developmental hematopoietic stem cell variation explains clonal hematopoiesis later in life. Nat. Commun. 2024;15(1):10268.

36. Gillis N, Padron E, Wang T, et al. Pilot Study of Donor-Engrafted Clonal Hematopoiesis Evolution and Clinical Outcomes in Allogeneic Hematopoietic Cell Transplantation Recipients Using a National Registry. Transplant. Cell. Ther. 2023;29(10):640.e1–640.e8.

37. Jain P, Duddu AS, Jolly MK. Stochastic population dynamics of cancer stemness and adaptive response to therapies. Essays Biochem. 2022;66(4):387–398.

38. Bukkuri A. Modeling stress-induced responses: plasticity in continuous state space and gradual clonal evolution. Theory Biosci. 2024;143(1):63–77.

39. Vipparthi K, Hari K, Chakraborty P, et al. Emergence of hybrid states of stem-like cancer cells correlates with poor prognosis in oral cancer. iScience. 2022;25(5):104317.

40. Su Y, Wei W, Robert L, et al. Single-cell analysis resolves the cell state transition and signaling dynamics associated with melanoma drug-induced resistance. Proc. Natl. Acad. Sci. 2017;114(52):13679–13684.

41. Mohammadi F, Visagan S, Gross SM, et al. A lineage tree-based hidden Markov model quantifies cellular heterogeneity and plasticity. Commun. Biol. 2022;5(1):1258.

42. Burkhardt DB, Juan BPS, Lock JG, Krishnaswamy S, Chaffer CL. Mapping Phenotypic Plasticity upon the Cancer Cell State Landscape Using Manifold Learning. Cancer Discov. 2022;12(8):1847– 1859.

43. Cho H, Ayers K, Pills L de, et al. Modelling acute myeloid leukaemia in a continuum of differentiation states. Lett. Biomath. 2018;5(sup1):S69–S98.

44. Ooi QX, Plan E, Bergstrand M. A tutorial on pharmacometric Markov models. CPT: Pharmacomet. Syst. Pharmacol. 2024;

45. Rockne RC, Branciamore S, Qi J, et al. State-Transition Analysis of Time-Sequential Gene Expression Identifies Critical Points That Predict Development of Acute Myeloid Leukemia. Cancer Res. 2020;80(15):3157–3169.

46. Frankhouser DE, Rockne RC, Uechi L, et al. State-transition modeling of blood transcriptome predicts disease evolution and treatment response in chronic myeloid leukemia. Leukemia. 2024;38(4):769–780.

47. Martínez-Rubio Á, Chulián S, Niño-López A, et al. Computational flow cytometry immunophenotyping at diagnosis is unable to predict relapse in childhood B-cell Acute Lymphoblastic Leukemia. Comput. Biol. Med. 2025;188:109831.

48. Stolpa W, Mizia-Malarz A, Zapała M, Zwiernik B. Can CD34+CD38− lymphoblasts, as likely leukemia stem cells, be a prognostic factor in B-cell precursor acute lymphoblastic leukemia in children? Front. Pediatr. 2023;11:1213009.

49. Newton PK, Mason J, Bethel K, et al. A Stochastic Markov Chain Model to Describe Lung Cancer Growth and Metastasis. PLoS ONE. 2012;7(4):e34637.

50. Newton PK, Mason J, Bethel K, et al. Spreaders and Sponges Define Metastasis in Lung Cancer: A Markov Chain Monte Carlo Mathematical Model. Cancer Res. 2013;73(9):2760–2769.

